# Interoceptive Accuracy Modulates Electroencephalogram During Music Recall Tasks

**DOI:** 10.1101/2025.05.09.652994

**Authors:** Kazuki Matsunaga, Ingon Chanpornpakdi, Toshihisa Tanaka

## Abstract

The ability to perceive and recall music varies with individuals, depending on musical experience, age, and emotional memory. Emotional memory processing occurs in the insular cortex, which is also implicated in interoception, the perception of internal bodily states, suggesting a potential link to music recall. However, the direct relationship between interoception and music recall remains largely unexplored. We hypothesize that individual differences in music recall are influenced by interoceptive accuracy. To test this, we conducted an electroencephalogram (EEG) experiment where participants listened to and recalled both familiar and unfamiliar music. Participants’ interoceptive accuracy was assessed through a heartbeat counting task. We observed greater alpha power suppression during recall periods in familiar music than in unfamiliar music. Furthermore, participants with higher interoceptive accuracy exhibited stronger alpha power suppression during music recall. These findings suggest that music recall involves interoceptive attention. Considering the insular cortex’s role in both interoception and emotional memory, it may play a critical role in the neural processes underlying music recall, which should be further investigated.

## Introduction

Music has deep-rooted origins in human cultures and it is a fundamental component of daily life. [1]. Humans can produce music using instruments or their voices, listen to it, or recall it internally without the actual sound [2]. These activities serve as entertainment for humans and also influence emotions, cognition, and memory [3–5]. Recalling music plays a crucial role in these activities, enabling humans to employ memory for emotional and cognitive functions. For instance, recalling a certain melody or rhythm has been shown to vividly revive past experiences or events [5]. Various factors, such as musical experience and age, can influence an individual’s ability to perceive and recall music. For example, musicians have a better perception of pitch changes in melodies than non-musicians [6], and harmonic perceptual functions decline with aging [7]. These factors are also believed to influence music recall [8, 9] and have been widely studied through behavioral experiments and brain imaging.

Apart from those factors, another internal characteristic that influences the perception and recall of music is interoception, which is the ability to sense internal bodily states, such as one’s heartbeat, breathing, and emotional changes [10]. It is primarily processed in the insular cortex in the brain [11], which is also involved in emotion recall from memory [12]. Furthermore, an increase in insular cortex activity was observed as the emotion changed during music listening [13], suggesting a relationship between music listening and insular cortex function [14]. Regarding the relationship between interoception and music perception, it has been reported that interoceptive accuracy (IAcc) [15], the ability to accurately perceive internal physiological signals, serves a crucial function. The individuals with lower IAcc perform better at perceiving rhythms than those with higher IAcc [16], and show less emotional change during music listening [17]. In addition, the study in IAcc and word recall shows that the participants with higher IAcc demonstrated better memory processing for words [18]. Given the similarities between music and language processing in the brain [19], it is suggested that participants with stronger interoceptive accuracy may exhibit better memory processing during music recall.

Music exhibits a temporal variation; therefore, imaging techniques with high temporal resolution are essential to capture the neural changes due to music. To measure the music-related neural responses, electroencephalography (EEG) or magnetoencephalography (MEG) is commonly used, as they offer high temporal resolution. Typical EEG responses during the music listening include event-related potentials (ERPs) [20], steady state-evoked potentials (SSEPs) [21], and power modulations in specific frequency bands [22]. Representative ERPs during music listening are N100, which appears as a negative peak around 100 ms after stimulus onset, and P200, which appears as a positive peak around 200 ms after stimulus onset [23]. SSEPs are related to the steady tempo of the music, producing clear peaks in the frequency spectrum at integer multiples of the tempo [24]. Regarding the power modulations, it has been reported that alpha (8–12 Hz) and low beta (12–16 Hz) power are more suppressed when listening to familiar music compared to unfamiliar music [22]. Furthermore, Müller et al. [25] used MEG to observe alpha power modulations during music recall. They asked participants to listen to musical stimuli in which a portion of the music was replaced by pink noise, and then asked them to recall the missing segment of the music. Time-frequency analysis of MEG signals during this task revealed that the alpha power was more suppressed when participants imagined a familiar music piece, compared to when they imagined an unfamiliar music piece [25]. Additionally, the alpha power modulation has been linked to the shifts in attention, particularly when focusing on internal (interoceptive) signals [26, 27]. Notably, interoceptive accuracy has been shown to be associated with various cognitive processes, including word recall [18], emotion recall [15], emotional changes induced by music [13], and rhythm perception [16]. Consequently, interoceptive accuracy may also contribute to alpha power suppression during music recall and play an essential role in music recall. However, limited studies have directly examined the influence of interoceptive accuracy on music recall itself, despite its established connections to related domains.

Therefore, in this paper, we aim to clarify the effect of interoceptive accuracy on EEG signals during music recall by proposing the following two hypotheses:

1. EEG responses differ between situations in which music is recalled and situations in which it is not.
2. When music is recalled, EEG responses differ between individuals with high and low interoceptive accuracy.

To test these hypotheses, we conducted an experiment in which participants performed a music recall task and a task assessing interoceptive accuracy.

## Materials and methods

### Participants

Twenty-two participants (18 males, 4 females, 20 right-handed, 2 left-handed, mean age = 23.9 ± 6.06 years, range = 20–49 years) were volunteered for the experiment. They had no history of neurological diseases (including epilepsy), mental disorders, or hearing impairments. Six participants had received formal musical education (7.50 ± 7.01 years, range = 1–20 years). The informed consent was obtained from all participants in accordance with the ethical approval by the Research Ethics Committee of Tokyo University of Agriculture and Technology (Approval No. 240210-0599) before the experiment. No participants were recruited by faculty recommendation, nor did they receive academic credit for participation. Hereafter, the participants are denoted by ID: mk1p–mk17p and mk18–mk22.

### Experimental Environment

To prevent external noise interference, the experiment was conducted in a soundproof room with the environment. The auditory stimulus was presented through a loudspeaker (NS-F210, YAMAHA, Japan), and instructions were given to participants via a display. The display (GL2750HM, BenQ, Taiwan) was positioned 1 m away from the participant. The participant’s head was fixed on a chinrest to prevent head movement during the experiment.

### Musical Stimuli

We used the melodies employed in the study by Ito et al. [28] on music listening and recall as the stimulus. A total of 40 melodies were used (listed in Supplementary Table S1 for all the musical pieces). The list consisted of 20 melodies that were generally considered familiar and 20 melodies that were considered unfamiliar, selected based on research on familiarity with music [20, 21]. The tempo of all melodies was standardized to 2.5 Hz [21, 28]. The musical score of the melodies was electronically produced with piano sounds in a musical instrument digital interface (MIDI) format. Three non-overlapping 12.8 s (8-bar) segments were extracted from each melody. Additionally, six notes (from the first note of bar 6 to the second note of bar 7) in each melody were replaced with rests, creating a 2.4 s silent interval to induce mental imagery of the melody without the actual sound. An example score of the melody segments used as a stimulus is shown in Fig. 1. In this manner, a total set of 120 stimuli was constructed (40 pieces × 3 segments). Each MIDI-based musical stimulus was saved as a WAV file at a sampling rate of 44,100 Hz and presented during the experiment.

**Fig 1.**
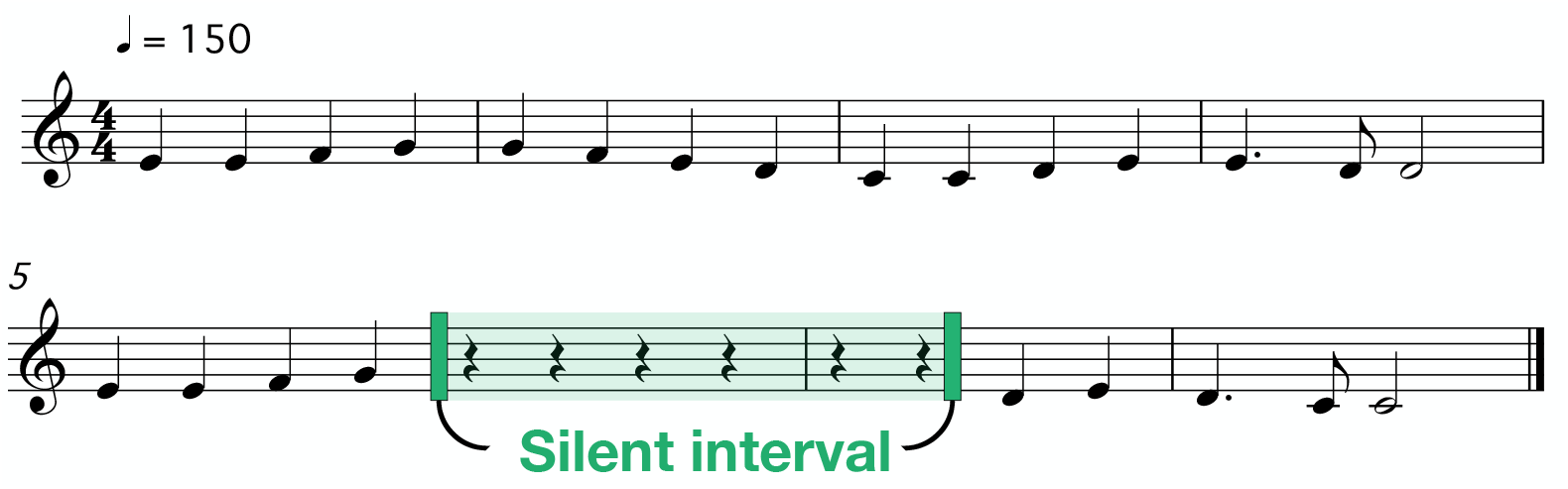
An example of musical stimulus in a single trial

### Experimental Tasks

#### Music Recall Task

While listening to the melody, participants were instructed to imagine the melody internally as it played. They were instructed to keep imagining the silent portion of the melody during the silent interval. Participants were informed that a silent interval would appear somewhere in the melody but were not told the exact timing. During the task, participants fixated on a cross presented at the center of the display. After listening to a melody containing a silent interval, participants were asked to rate their familiarity with the melody. The underlying idea behind this experimental design was that familiar music is more easily recalled, whereas recalling unfamiliar music may not be as easy [29]. We categorized music pieces to which each participant listened into the familiar and unfamiliar groups.

Familiarity ratings were collected on a 5-point Likert scale (5: extremely familiar, 4: moderately familiar, 3: somewhat familiar, 2: slightly familiar, 1: not at all familiar) [30]. Participants were instructed as follows:

- If they knew the melody and if the melody following the silent interval matched their imagined continuation, they were to select “5.”
- If they knew the melody but the post-silence melody differed from what they had imagined, they were to select “4.”
- If they were unsure or only recalled the melody after listening, they were instructed to select “3.”
- If they knew it but could not imagine it, they were to select “2.”
- If they did not know the melody at all, they were to select “1”.

The familiarity rating was done by clicking the corresponding on-screen option with a mouse cursor.

#### Heartbeat Counting Task

To quantify the accuracy of interoception, participants were asked to perform the heartbeat counting task (HCT) [31], in which they counted their heartbeats in their minds. They were instructed to refrain from estimating the number of heartbeats based on their resting heart rate or manually checking their pulse (e.g., on the wrist or neck) [32]. During the task, participants were also instructed to keep their gaze on a fixation cross at the center of the display. They silently counted their heartbeats for a set period and then reported the counted number via keyboard input. The duration of the counting period was not informed to the participants.

### Experimental Flow

The music recall task and HCT were administered to the participants. The music recall task was conducted first to measure EEG during music recall (listening to music while imagining it internally), followed by the HCT was conducted to assess the accuracy of interoception. EEG, EOG, and PPG were recorded throughout all tasks.

#### Music Recall Task Procedure

The flow of one trial is shown in Fig. 2. Each trial started with a 4-s waiting period, followed by the presentation of a 0.1 s beep sound (440 Hz), 12.8 s of musical stimulus, and a 2.2-s waiting period, as shown in Fig. 2a. After this waiting period, the melody familiarity rating screen was provided to the participants for them to report how familiar they were with the presented melody, as shown in Fig. 2b. The experiment consisted of 120 trials, each of which included the aforementioned steps. Twelve stimuli were presented in a block, with a total of 10 blocks and a short break provided after each block. The order of stimulus presentation was randomized. Prior to the main session, a practice session was conducted to ensure that participants understood the task. This practice session consisted of three trials which had the same structure as the main session with different melodies. The stimulus presentation program was created using C# 12.0.

**Fig 2.**
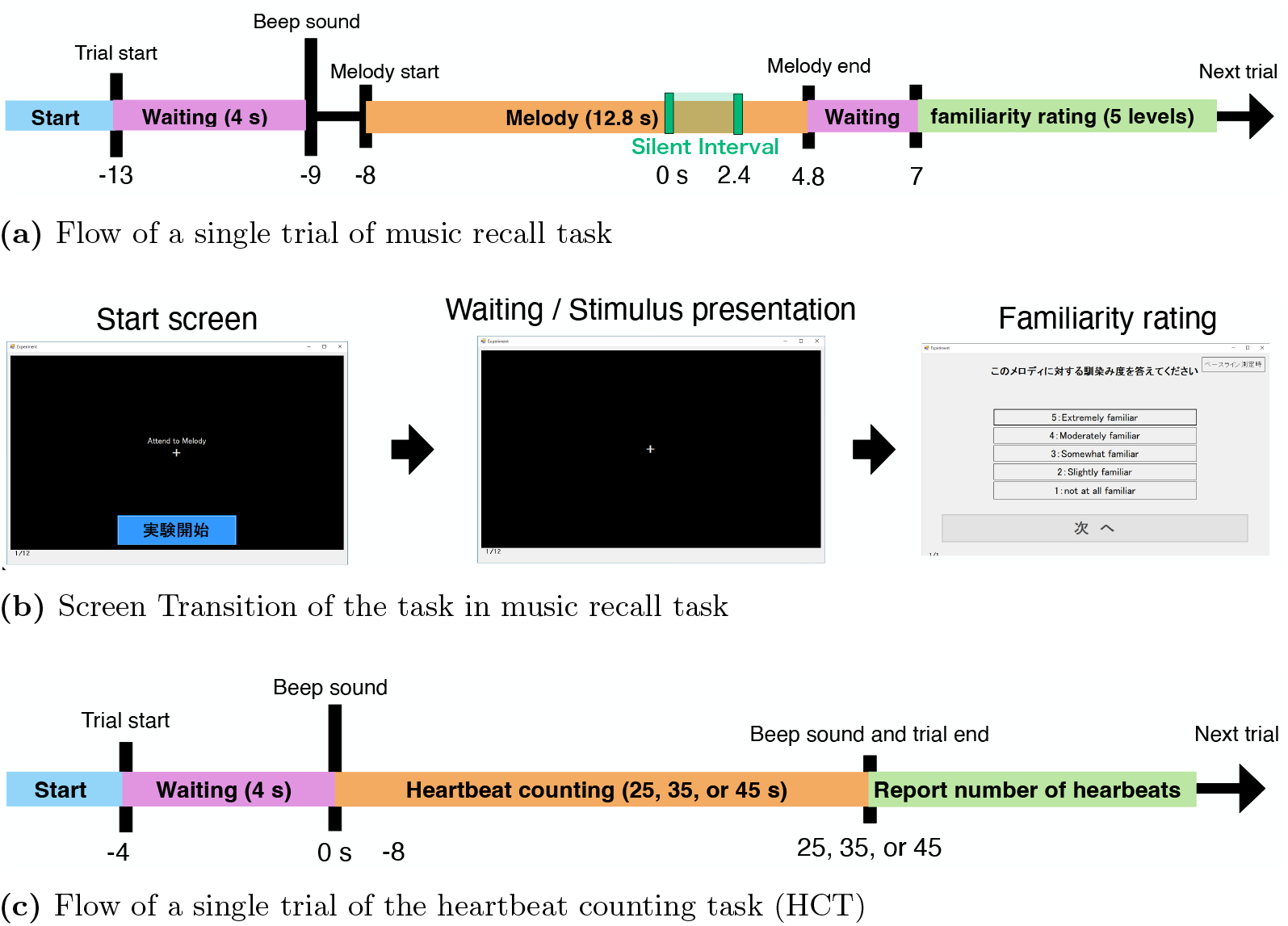
Experimental flow of a single trial in music recall and HCT task

#### Heartbeat Counting Task Procedure

The flow of one trial is shown in Fig. 2c. Similar to the music recall task, the trials started with a 4-s waiting period after the start of the trial. After the waiting period, two beep sounds were played. The first 0.1-s beep sound indicated the start of the counting period, and the subsequent 0.1-s beep sound indicated the end of the counting period. The participants were instructed to internally count their heartbeats and stop when a beep sound was played for the second time. After the counting period, participants reported the number of heartbeats they had counted. Three trials were conducted with the counting period of 25 s, 35 s, or 45 s in a randomized order [12, 31].

### Data Acquisition

This experiment recorded EEG, electrooculograms (EOG), and photoplethysmograms (PPG) as physiological signals. For EEG recording, 64 scalp electrodes mounted on a May 9, 2025 5/30 gel-based EEG head cap (TMSi, Twente Medical Systems International, Oldenzaal, The Netherlands) were used. The electrode arrangement used followed the international 10–10 system (Fp1, Fpz, Fp2, AF7, AF3, AF4, AF8, F7, F5, F3, F1, Fz, F2, F4, F6, F8, FT7, FC5, FC3, FC1, FCz, FC2, FC4, FC6, FT8, M1, T3, C5, C3, C1, Cz, C2, C4, C6, T4, M2, TP7, CP5, CP3, CP1, CPz, CP2, CP4, CP6, TP8, T5, P5, P3, P1, Pz, P2, P4, P6, T6, PO7, PO5, PO3, POz, PO4, PO6, PO8, O1, Oz, and O2). Additionally, EOG and PPG were recorded simultaneously with EEG to confirm blink, eye movements, and heart rate. EOG was measured using two bipolar electrodes (Microelectrodepair (bipolar), TMSi, The Netherlands). The two channels of EOG were horizontal EOG (HEOG) and vertical EOG (VEOG). The HEOG channel was placed near the outer canthus of the right eye, with a reference electrode placed near the outer canthus of the left eye. The VEOG channel was placed above the right eye with a reference electrode below the right eye. PPG was recorded using a finger clip sensor (8000AA, Nonin, USA) attached to the index finger of the left hand. All signals were amplified using TMSi Refa 72-channel amplifiers (TMSi, Twente Medical Systems International, Oldenzaal, The Netherlands)and digitized by the amplifier’s built-in ADC. The common average reference was used with the disposable ground electrode placed on the participant’s left wrist. The signals were recorded using Polybench (TMSi, Twente Medical Systems International, Oldenzaal, The Netherlands) software at a sampling rate of 2,048 Hz.

### EEG Preprocessing

EEG preprocessing was performed using Python 3.13.0 and MNE-Python 1.8.0 [33] according to the following steps:

1. A 50 Hz notch filter and a 1–40 Hz FIR band-pass filter were applied to remove electrical noise and other artifacts.
2. The recorded EEG signal was downsampled to 256 Hz.
3. The EEG was re-referenced to the average potential of the M1 and M2 electrodes placed on the mastoids.
4. Independent component analysis (ICA) was performed to decompose the EEG into 20 components using the extended infomax algorithm [34] (implemented using `mne.preprocessing.ICA`). The components that had a Pearson correlation coefficient with the EOG signal of 0.9 or higher were then eliminated, resulting in an average component removal of 2.42 ± 1.12 components per participant.
5. The EEG was segmented into epochs for each trial. The time range for each epoch was set from [− 13, 7] s, with 0 s corresponding to the onset of the silent interval. Thus, each epoch covered the 20 s from the trial start to the beginning of the familiarity rating.
6. The epochs with the peak-to-peak amplitude exceeding 200 *μ*V were removed to further eliminate any remaining eye or muscle artifacts that could not be captured by ICA. An average of 18.1 ± 19.4 epochs were excluded per participant.

### EEG Epoch Labeling

Based on the familiarity ratings for each melody, the EEG epochs were divided into two groups [21]. If the rating was “5” or “4”, the epoch was labeled “familiar” and if the rating was “2” or “1”, the epoch was labeled “unfamiliar.” Epochs labeled “familiar” were treated as instances where the participant could imagine the melody, whereas epochs labeled “unfamiliar” were treated as instances where the participant could not May 9, 2025 6/30 imagine the melody. Epochs rated “3” were excluded from the analysis. On average, 3.09 ± 3.95 epochs per participant were excluded for being rated as “3.”

### HCT Score Calculation

Interoceptive accuracy (IAcc), the score from the HCT, was computed using the following standard formula [31, 32]:

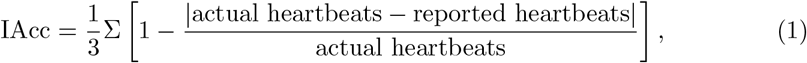

where actual heartbeats is the participant’s actual number of heartbeats during the task (derived from PPG) and reported heartbeats is the number the participant counted and reported during the task. The range of IAcc is from 0 to 1, with the higher values indicating greater interoceptive accuracy.

### Comparison Between Familiarity Groups

#### Time-Frequency Analysis

A time-frequency analysis was conducted across all participants to compare changes in EEG power between the familiar and unfamiliar groups during music recall. First, we computed the relative spectrogram concerning the baseline for each epoch after preprocessing using a log ratio:

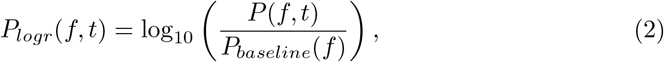

where *t* is the time in seconds, with 0 s defined as the onset of the silent interval, *P* (*f, t*) is the spectrogram of the epoch, and *P*_*baseline*_(*f*) is the power spectrum averaged over the baseline period [− 11, − 9]s. The spectrogram was estimated using the multi-taper method [35, 36], implemented by `mne.time frequency.tfr multitaper` in MNE-Python.

Next, the difference in the relative spectrogram between familiar and unfamiliar was computed as

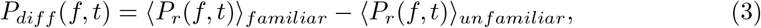

where ⟨*P*_*r*_(*f, t*)⟩ _*familiar*_ and ⟨*P*_*r*_(*f, t*)⟩ _*unfamiliar*_ are the epoch-averaged relative spectrograms for the familiar and unfamiliar groups, respectively. Finally, a cluster-based permutation test [37] was performed on *P*_*diff*_ (*f, t*) for each electrode, taking multiple comparisons into account. The `mne.stats.permutation_cluster_test` function was used with 10,000 permutations and a significance level of 0.05.

#### ERP Analysis

To examine event-related potentials (ERPs) elicited by the stimulus, we first computed the averaged EEG waveform for each participant and each electrode under familiar and unfamiliar groups. The baseline was the rest period [− 11, − 9]s. We checked for N100, P200, P300, and late positive potential (LPP) components around the onset of the silent interval. Specifically, we examined the mean amplitude in the following time windows: N100 in [0.09, 0.12]s, P200 in [0.17, 0.24]s, P300 in [0.32, 0.38]s, and LPP in [0.4, 0.8]s [38, 39]. For each ERP time window, a two-sided one-sample t-test against zero was conducted for each group and electrode. The significance level was set to 0.05, and the p-values were corrected for multiple comparisons using the Holm-Bonferroni method.

We also examined N100, P200, and P300 elicited by the resumption of the musical stimulus after the silent interval. The time windows for these components were: N100 in [2.49, 2.52]s, P200 in [2.57, 2.64]s, and P300 in [2.72, 2.78]s. LPP was not tested for the post-silence onset due to the 2.5 Hz stimulus tempo, which introduces the next note onset 0.4 s after the previous note.

#### SSEP Analysis

To compare the frequency characteristics of EEG between melody-present and silent intervals, we computed the steady state-evoked potential (SSEP) for both the familiar and unfamiliar groups. Epochs were subdivided into two segments: the melody interval [− 5, 0] s and the silent interval [0, 2.4] s. The baseline period was [− 11, − 9] s. For each group (familiar/unfamiliar) and each interval (melody/silent), the epoch-averaged EEG was calculated for each electrode and then underwent a Fourier transform (via `numpy.fft.rfft`) to estimate its power spectrum. Finally, the power spectra were averaged across all electrodes.

From the obtained electrode-averaged power spectrum, we extracted the peak amplitudes at 2.5 Hz, 5 Hz, 7.5 Hz, and 10 Hz. These peaks correspond to the tempo of the musical stimulus and its harmonics [24]. A two-way repeated-measures ANOVA was performed on each peak amplitude. The independent variables were familiarity (familiar/unfamiliar) and interval (melody/silent), and the dependent variable was the peak amplitude for each participant. A significance level of 0.05 was used. Subsequently, paired-sample two-sided t-tests were conducted for each familiarity group (familiar/unfamiliar) to compare the peak amplitude between the melody and silent intervals as a post-hoc test for the two-way ANOVA. Additionally, for each interval (melody/silent), a paired t-test was conducted to compare the peak amplitude between the familiar and unfamiliar groups. The significance level was set to 0.05, and p-values were corrected using the Holm-Bonferroni method.

#### Comparison of Spectrograms during Music Recall Between Interoception Groups

To assess the effect of interoception on EEG during music recall, participants were grouped based on high and low interoceptive accuracy. A time-frequency analysis was then performed to compare the changes in EEG power across different familiarity groups and interoceptive groups.

#### Selection of Target Participants Based on IAcc

All participants were sorted by their IAcc scores, and those scores above the third quartile were classified into the high-interoceptive accuracy group (high IAcc). On the other hand, those whose scores were below the first quartile were classified into the low-interoceptive accuracy group (low IAcc). Each group contained 6 participants, and the remaining 10 participants were excluded from further analyses. A one-sided independent t-test was conducted to compare IAcc between the high IAcc and low IAcc groups, with the alternative hypothesis that the high IAcc group’s IAcc is significantly higher than that of the low IAcc group. The significance level was set at 0.05.

#### Comparison of Spectrograms during Music Recall

The relative spectrograms for each familiarity group (familiar, unfamiliar) to the baseline were computed as described in time-frequency analysis. This process yielded four groups (familiar group for high IAcc participant group; high IAcc familiar, unfamiliar group for high IAcc participant group; high IAcc unfamiliar, familiar group for low IAcc participant group; low IAcc familiar, and unfamiliar group for low Iacc participant group; low IAcc unfamiliar). For each of these four groups, we averaged the spectrograms across epochs and electrodes, and then extracted the mean alpha band power ([8, 12] Hz) and low-beta band power ([13, 16] Hz) [22] from two time segments: an early silent segment [0, 0.05]s and a late silent segment [2, 2.05]s.

The mean values were then subjected to a mixed-design two-way ANOVA with familiarity (familiar/unfamiliar) and interoception group (high IAcc/low IAcc) as independent variables, and each participant’s mean power as the dependent variable. The significance level was set to 0.05, and tests were conducted separately for each time segment and frequency band. Sequentially, t-tests were performed as posthoc tests. Within each interoception group (high IAcc or low IAcc), a two-sided paired t-test was used to compare the power between the familiar and unfamiliar groups for each time segment and frequency band. Additionally, for each familiarity group (familiar or unfamiliar), a two-sided independent t-test was performed to compare the power between the high IAcc and low IAcc groups. The significance level was set at 0.05, and p-values were corrected using Tukey’s method for multiple comparisons.

Finally, to evaluate topographical differences between interoception groups during music recall (familiar group), we computed

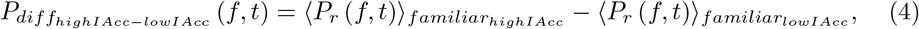

where 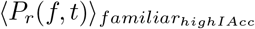 and 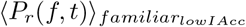 denote the epoch-averaged relative spectrograms under the familiar group for the high IAcc and low IAcc groups, respectively. A cluster-based permutation test was then conducted on 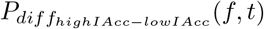 for each electrode, following the procedure in time-frequency analysis, with a significance level of 0.05.

## Results

### HCT Scores for Each Participant

The heartbeat counting task (HCT) scores, denoted as IAcc for each participant, showed an average of 0.811 ± 0.136, with a minimum of 0.541 and a maximum of 0.990 (refer to Supplementary Table S2 for IAcc of each participant).

### Comparison Between Familiarity Groups in Music Recall Task

#### Time-Frequency Analysis for Familiarity

The sample of spectrograms during music listening and recall for each familiarity group (familiar and unfamiliar groups) is shown in Fig. 3a. The six electrodes were selected from 64 (left frontal: F3, right frontal: F4, left central: C5, right central: C6, left parietal: P3, right parietal: P4) [28]. Both familiar and unfamiliar groups showed a tendency for suppression of alpha and beta band spectrograms compared to the resting state, as shown in Fig. 3a. Furthermore, during music listening, an increase in spectrograms below the theta band was observed in both groups and immediately after music resumption.

**Fig 3.**
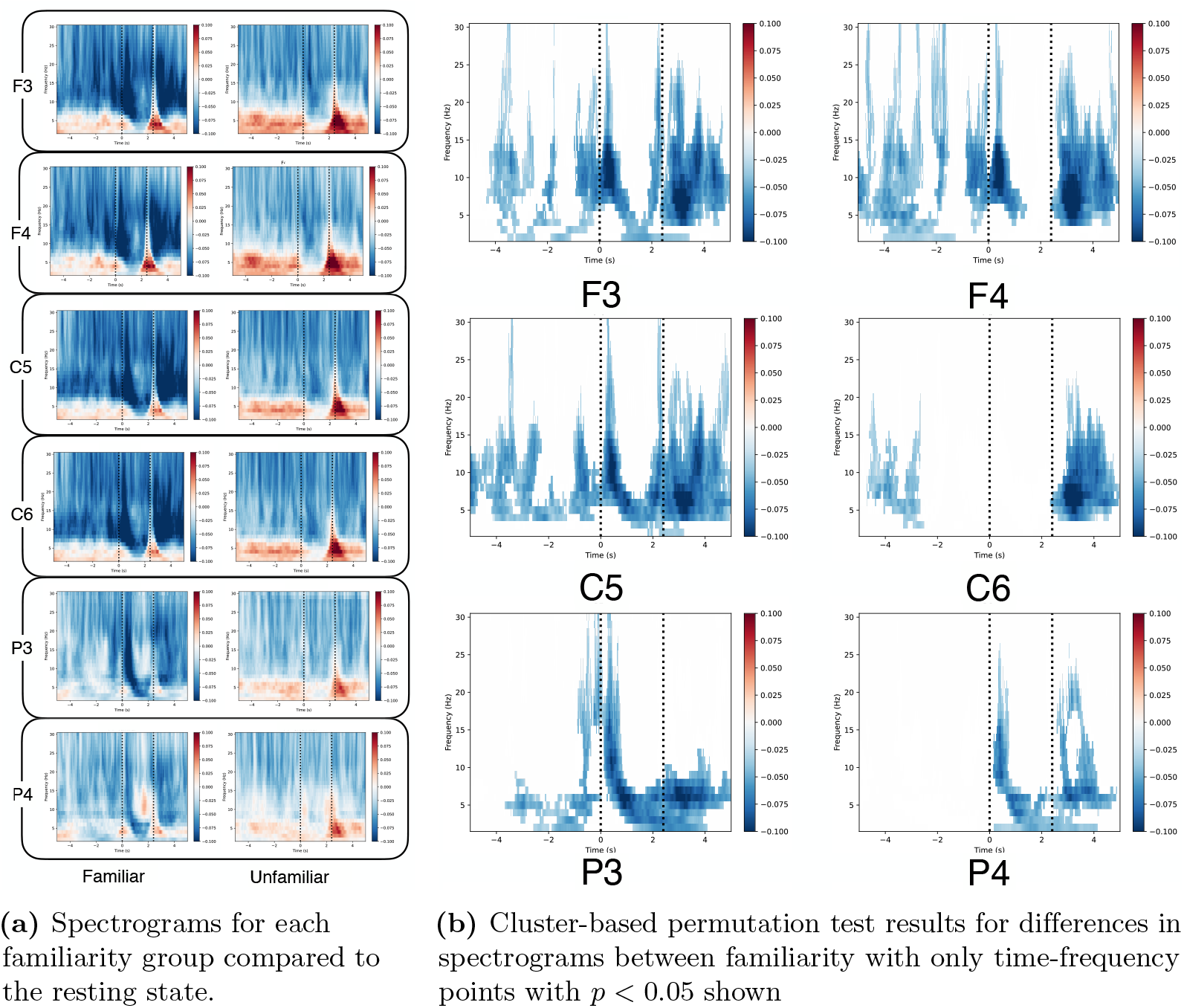
Excerpt results of the time-frequency analysis at the selected electrodes (left frontal: F3, right frontal: F4, left central: C5, right central: C6, left parietal: P3, right parietal: P4). The dashed line at 0 s marks the onset of the silent interval, and the dashed line at 2.4 s marks the end of the silent interval.

In addition, we performed the cluster-based permutation test for differences in spectrograms between the two groups. The partial results of the cluster-based permutation test are shown in Fig. 3b, based on the selected electrodes in Fig. 3a. The results show that the alpha and beta suppression were significantly stronger when the familiar melody stimuli were presented. In contrast, no significant difference was observed in the increase in the theta band when comparing the familiarity groups. The differences in spectrograms between the familiarity groups for each of the 64 electrodes, arranged according to electrode layout, are shown in Supplementary Fig. S3a, and the significant clusters identified by the cluster-based permutation test for each electrode are presented in Supplementary Fig. S3b. In most electrodes, the familiar group exhibited significantly lower spectrograms during music listening and imagery than the unfamiliar group. However, some electrodes in the right central and right parietal areas showed no significant differences between familiarity groups during music listening and silent periods.

#### ERPs at Silent Interval

The grand-averaged ERP across all the electrodes in familiar and unfamiliar groups is shown in Fig. 4a. From the figure, a small negative peak N100-like ERP component could be observed only in the familiar group. The electrode-wise ERPs in the familiar group aligned to the onset of the silent period are shown in Supplementary Fig. S4a, and those in the unfamiliar group are shown in Supplementary Fig. S4b, arranged according to electrode layout. Considering the ERP for each electrode, a small positive peak appeared after the onset of the silent period, mainly in parietal and occipital electrodes in the familiar group (See Supplementary Fig. S4a for more details). In contrast, no such peak was observed in the unfamiliar group after the silent period onset (See Supplementary Fig. S4b for more details).

**Fig 4.**
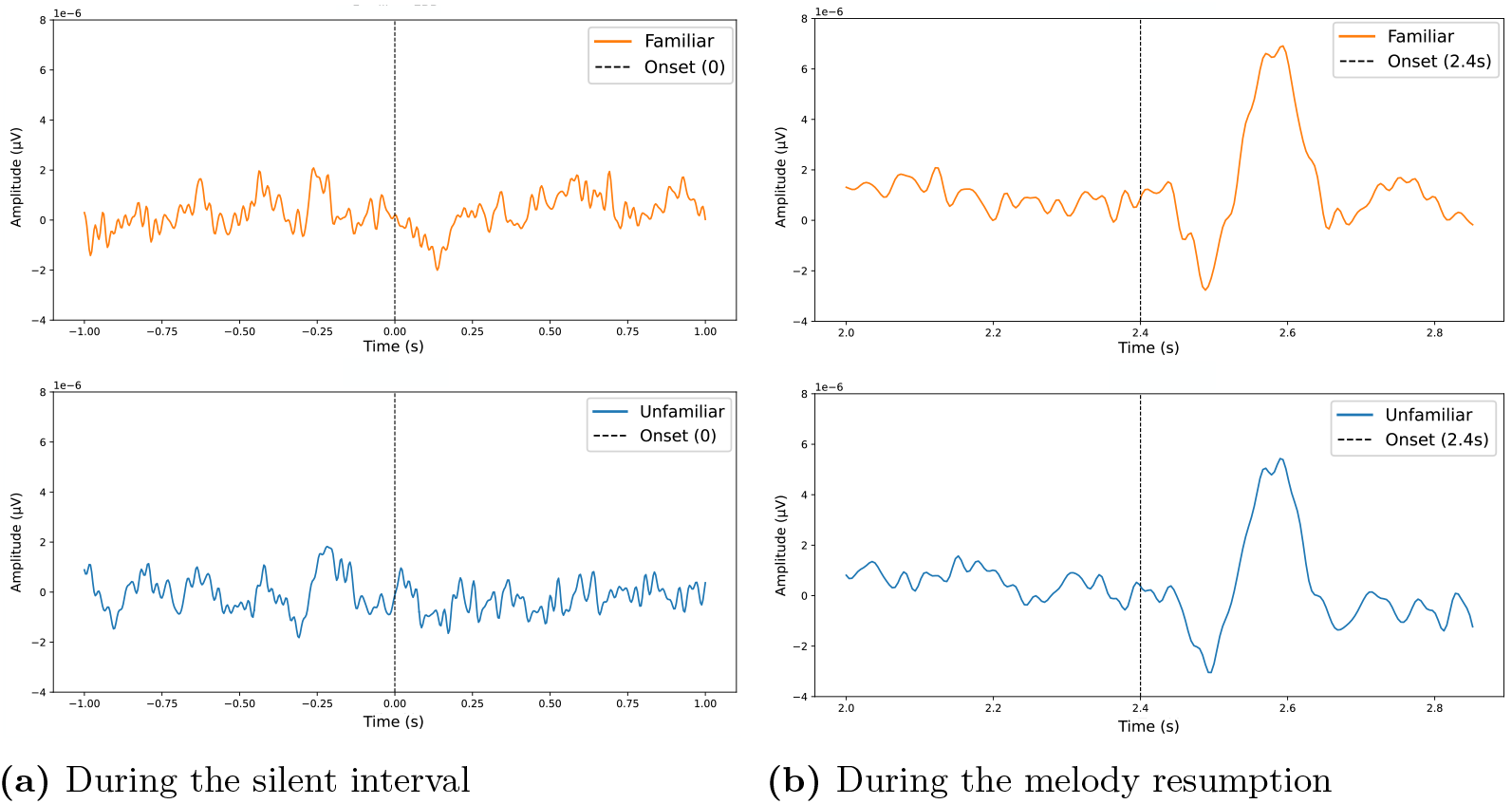
Grand-averaged ERP across all electrodes during familiar and unfamiliar groups. The orange line represents the familiar group, and the blue line represents the unfamiliar group. The dotted black line shows the onset time of the corresponding interval.

One-sample *t*-tests were conducted for each familiarity group and ERP component May 9, 2025 10/30 and revealed that for the N100 component, significant differences from zero were observed at right hemisphere electrodes (F4, F6, FC6, C6, FT8, T4, TP8, CP6) in the familiar group (see Supplementary Table S5 for more details), and at occipital electrodes (Oz, O2, POz, PO4) in the unfamiliar group (see Supplementary Table S6 for more details). For the P200 and P300 components, significant differences for the P200 component were observed at C6, T4, TP8, CP6 (see Supplementary Table S7 for more details) and for P300 components at CP6, P3, Pz, P4, T6, POz, P5, P1, P2, P6, PO3, PO4, PO6, TP8, PO7, PO8 (see Supplementary Table S9 for more details) in the familiar group. No significant differences were found in any electrodes in the unfamiliar group for P200 and P300 components (see Supplementary Tables S8 and S10 for more details, respectively). Regarding the LPP component, significant differences from zero were observed mainly in parietal and occipital regions in the familiar group, specifically at CP5, CP1, CP2, T5, P3, Pz, P4, POz, O1, Oz, O2, CPz, P5, P1, P2, P6, PO5, PO3, PO4, PO6, PO7, and PO8 (see Supplementary Table S11 for more details). On the other hand, only PO3 showed a significant difference in the unfamiliar group (see Supplementary Table S12 for more details).

#### ERPs at Melody Resumption

The grand-averaged ERP across all electrodes is shown in Fig. 4b. From the figure, we could observe three ERP components, which are N100, P200, and small P300 peaks in the familiar group. Except for the P300 peak, we also observed similar ERP peaks that were N100 and P200 peaks in the unfamiliar group. The electrode-wise ERPs in response to the melody resumption following the silent period in the familiar group are shown in Supplementary Fig. S13a, and those in the unfamiliar group are shown in Supplementary Fig. S13b, arranged according to electrode layout. As shown in Fig. S13a and Fig. S13b, ERP-like peaks appeared mainly in the parietal and central regions in both familiarity groups.

One-sample *t*-tests conducted for each familiarity group and ERP component revealed that, for both groups, significant differences from zero were observed in the N100 and P200 components at most electrodes. For the N100 component, significant differences from zero were observed at all electrodes except for T6, M1, and M2 in the familiar group (see Supplementary Table S14 for more details), and at all electrodes except for M1 and M2 in the unfamiliar group (see Supplementary Table S15 for more details). Regarding the P200 component, significant differences from zero were found at all electrodes except for M1, M2, and occipital electrodes at O1, Oz, PO5, PO7, and PO8 in the familiar group (see Supplementary Table S16 for more details). Similarly, significant differences from zero were found at all electrodes except for M1, M2, O1, Oz, O2, PO1, PO2, PO3, PO4, PO5, PO6, PO7, and PO8 for the unfamiliar group (see Supplementary Table S17 for more details). On the other hand, no significant differences were found in the P300 component at any electrode in either group (see Supplementary Tables S18 and S19 for more details, respectively).

#### Comparison of SSEP by Familiarity and Interval

The EEG power spectral density averaged across participants and electrodes is shown in Fig. 5. Both familiarity groups exhibited spectral peaks at multiples of the melody tempo (2.5 Hz, 5.0 Hz, 7.5 Hz, and 10 Hz) during the melody interval, while no such peaks were observed during the silent interval. The results of a two-way repeated-measures ANOVA conducted on the power spectral density averaged across all electrodes, focusing on the multiples of the melody tempo (2.5 Hz, 5.0 Hz, 7.5 Hz, and 10 Hz) (see Supplementary Table S20). The significant main effects were found for the familiarity factor (2.5 Hz: *F* = 19.3, *p <* .001, *η*^2^ = 0.921; 5.0 Hz: *F* = 16.1, *p <* .001, *η*^2^ = 0.765; 7.5 Hz: *F* = 17.6, *p <* .001, *η*^2^ = 0.838) and for the interval factor (2.5 Hz: *F* = 76.2, *p* = .001, *η*^2^ = 3.63; 5.0 Hz: *F* = 176, *p <* .001, *η*^2^ = 8.38; 7.5 Hz: *F* = 57.4, *p <* .001, *η*^2^ = 2.73; 10 Hz: *F* = 43.7, *p <* .001, *η*^2^ = 2.08) (see Supplementary Table S20). For the familiarity factor, no significant difference was observed at 10 Hz (*F* = 0.311, *p* = 0.583, *η*^2^ = 0.0148). Additionally, a significant interaction was found only at 5.0 Hz (*F* = 17.2, *p* = .001, *η*^2^ = 0.818) and 7.5 Hz (*F* = 9.94, *p* = .01, *η*^2^ = 0.473).

**Fig 5.**
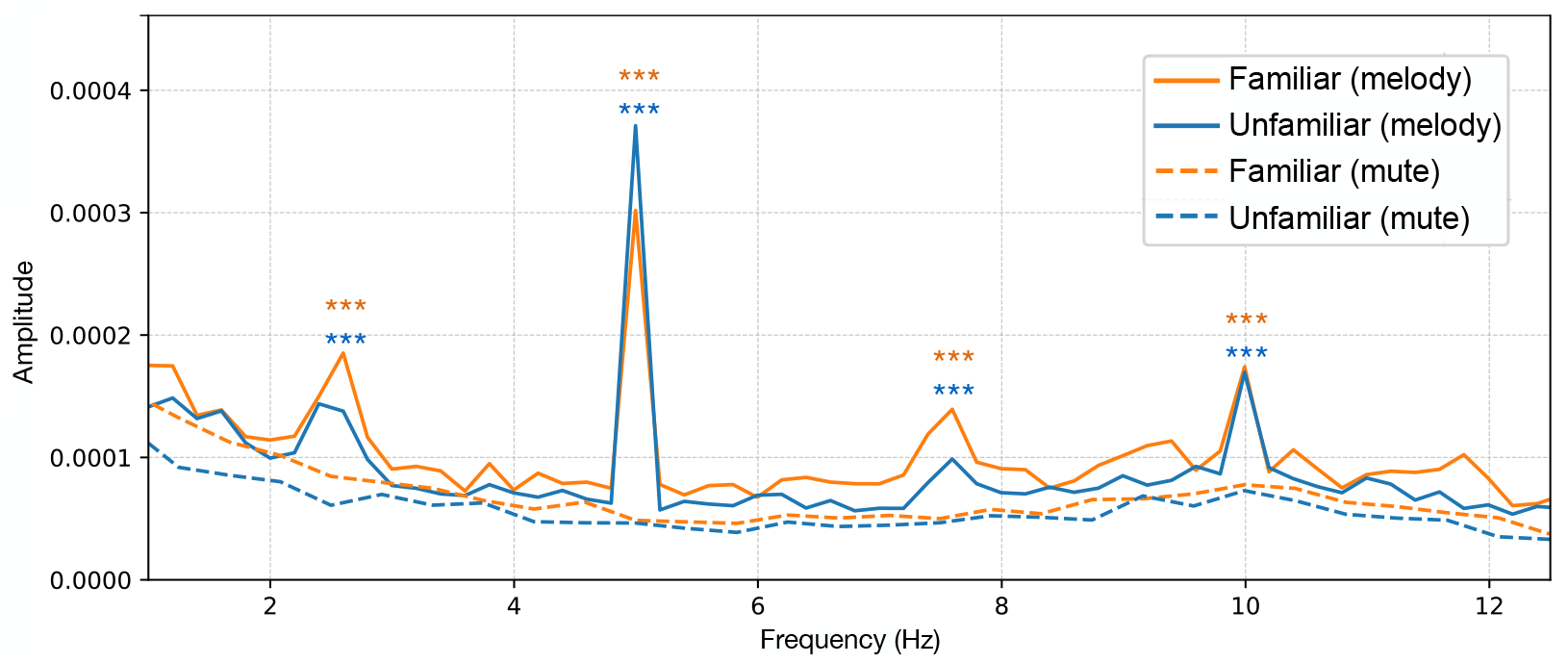
Power spectral density averaged across participants and electrodes. The orange solid line represents the melody interval for the familiar group, the orange dashed line represents the silent interval for the familiar group, the blue solid line represents the melody interval for the unfamiliar group, and the blue dashed line represents the silent interval for the unfamiliar group. \*\*\**p <* 0.001

The results of two-tailed paired *t*-tests, conducted as post-hoc analyses. During the melody interval, significant differences between familiarity groups were found at 2.5 Hz, 5.0 Hz, and 7.5 Hz (2.5 Hz: *t*(21) = 3.17, *p* = .0228, *d* = 0.677; 5.0 Hz: *t*(21) = − 4.36, *p <* .01, *d* = − 0.93; 7.5 Hz: *t*(21) = 3.96, *p <* .01, *d* = 0.844) (see Supplementary Table S21). In contrast, during the silent interval, a significant difference between familiarity groups was observed only at 2.5 Hz (*t*(21) = 3.58, *p* = .0105, *d* = 0.764). In addition, the comparisons between interval conditions revealed significant differences at all frequencies for both familiarity groups (see Supplementary Table S22).

### Between-Group Comparison During Music Recall

#### Comparison of IAcc Between Interoception Groups

Based on the IAcc scores, six participants were grouped into the high IAcc and low IAcc groups using the interquartile range (IQR) criterion. The participants assigned to the high IAcc group were participants 3, 4, 15, 16, 19, and 20, whereas those assigned to the low IAcc group were participants 5, 6, 7, 8, 9, and 12. The IAcc scores for each interoceptive group are shown as boxplots in Fig. 6, showing that the IAcc score for the high IAcc group was at 0.971 ± 0.02, while that for the low IAcc group was at 0.632 ± 0.06. An independent *t*-test revealed that the IAcc score in the high IAcc group was significantly higher than that in the low IAcc group (*t*(11) = 13.1, *p <* .001, *d* = 7.58).

**Fig 6.**
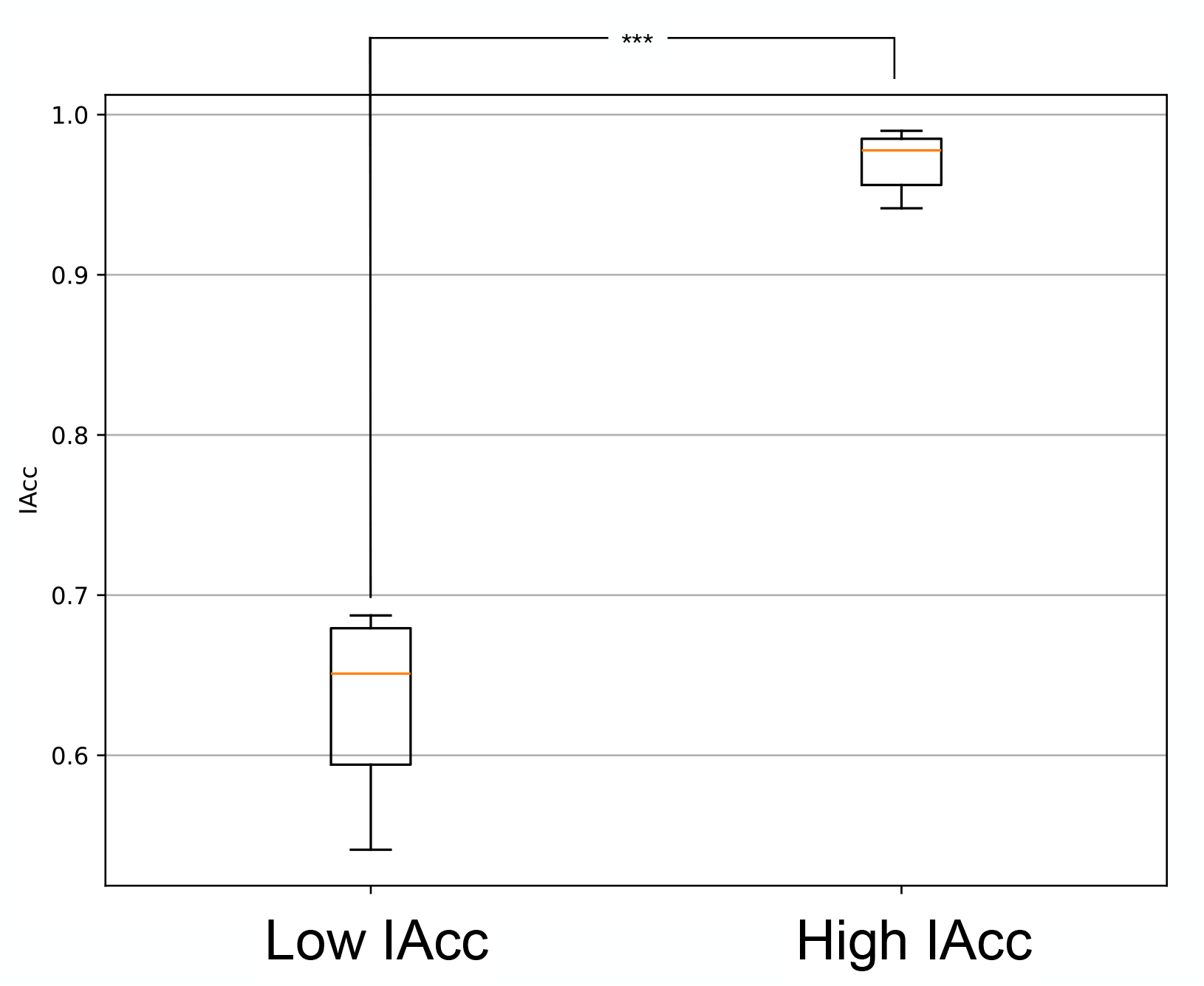
Boxplot of IAcc for the high IAcc low IAcc interoception groups (6 participants per group, * * * : *p <* .001)

#### Effect of Interoception on Spectrogram During Music Recall

The spectrograms with respect to baseline averaged across participants and electrodes for each familiarity and interoception group are shown in Fig. 7.

**Fig 7.**
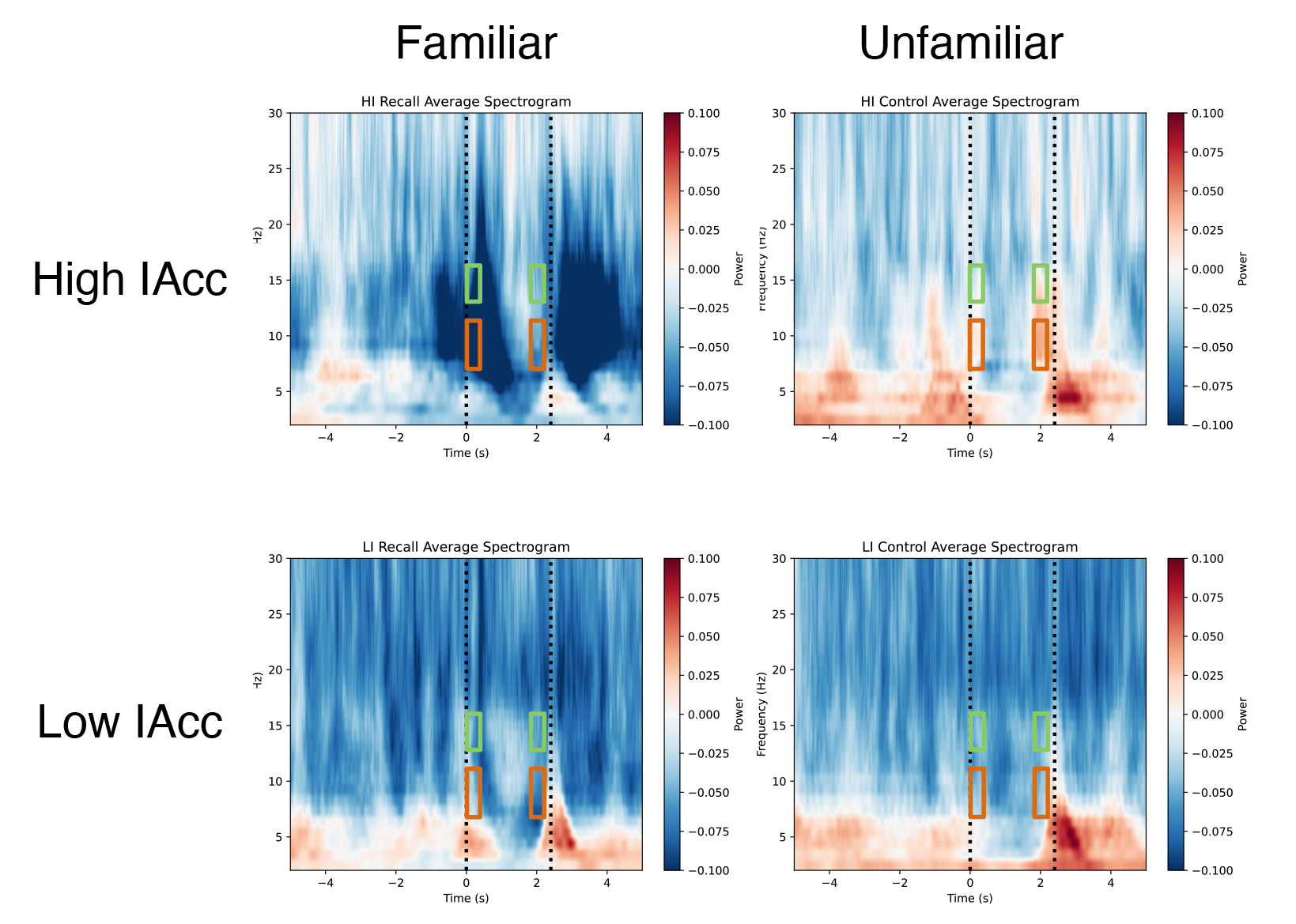
Grand-average spectrograms (across participants and electrodes) by familiarity and interoception group. The orange line indicates alpha power, and the green line indicates low-beta power. The dashed line at 0 s marks the onset of the silent interval, and the dashed line at 2.4 s marks the end of the silent interval.

The alpha and low-beta power during the silent interval ([0, 2.4] s) were strongly suppressed when compared to the resting state ([−12, − 9] s) in the high IAcc familiar group, as shown in Fig. 7. In contrast, the power suppression during the silent interval was less pronounced in the high IAcc unfamiliar group than in the high IAcc familiar group. Furthermore, in the low IAcc group, the difference in power between familiarity groups was small.

We performed a two-way mixed-design ANOVA on spectrograms averaged across all electrodes for alpha and low-beta power during the early silent interval ([0, 0.05] s) and those for the late silent interval ([2, 2.05] s). For alpha power during the onset of silence (early silent period), a significant main effect of the familiarity factor was found (*F* = 5.42, *p* = .042, 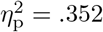), while no significant difference was found for low-beta power (see Supplementary Table S23). Moreover, significant interactions were found for both alpha and low-beta power (alpha: *F* = 11.75, *p* = .006, 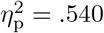; low-beta: *F* = 7.31, *p* = .022, 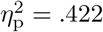). No significant main effect of the interoception group factor was found for either alpha or low-beta power. During the late silent interval, the trends toward significance for the main effect of the familiarity factor were observed on both alpha and low-beta power (alpha: *F* = 4.47, *p* = .061, 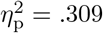; low-beta: *F* = 4.03, *p* = .072, 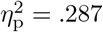), though these did not reach statistical significance (see Table S24). No significant main effects of the interoception group factor or interaction effects were found for either alpha or low-beta power.

We also performed two-tailed paired *t*-tests conducted as post-hoc analyses for the early and late silent intervals. During the onset of silence (early silent interval), a significant difference was observed only in the comparison between familiarity groups within the high IAcc group for alpha power (*t*(11) = − 4.071, *p* = .010, *d* = − 1.844) (see Supplementary Table S25). Additionally, a marginal trend was found in comparing interoception groups within the familiar group (*t*(11) = − 2.386, *p* = .117, *d* = − 1.378) based on the large Cohen’s d effect size. For low-beta power, no significant differences were found in any comparisons, though a marginal trend was observed for the comparison between familiarity groups within the high IAcc group (*t*(11) = − 3.037, *p* = .052, *d* = − 1.080). Regarding the late silent interval, a marginal trend was observed only for the comparison between familiarity groups within the high IAcc group for low-beta power (*t*(11) = − 2.593, *p* = .104, *d* = − 1.164) based on the Cohen’s d effect size (see Supplementary Table S26). In contrast, no significant differences or trends were found for other comparisons.

#### Comparison of Spectrograms by Electrode for Interoception Groups During Music Recall

The results of the cluster-based permutation test for differences between interoception groups (high IAcc, low IAcc) in the spectrograms of the familiar group are shown in Fig. 8, arranged according to the electrode layout. Regarding the spectrograms immediately after the onset of silence (early silence), the alpha power in the high IAcc group was significantly lower than that in the low IAcc group at central and right centro-parietal electrodes (FC2, C1, Cz, C2, CP6) as shown in Fig. 8. In contrast, the beta power in the high IAcc group was significantly higher than that in the low IAcc group at left temporal and occipital electrodes (C5, Pz, POz, PO5, PO7) during music listening.

**Fig 8.**
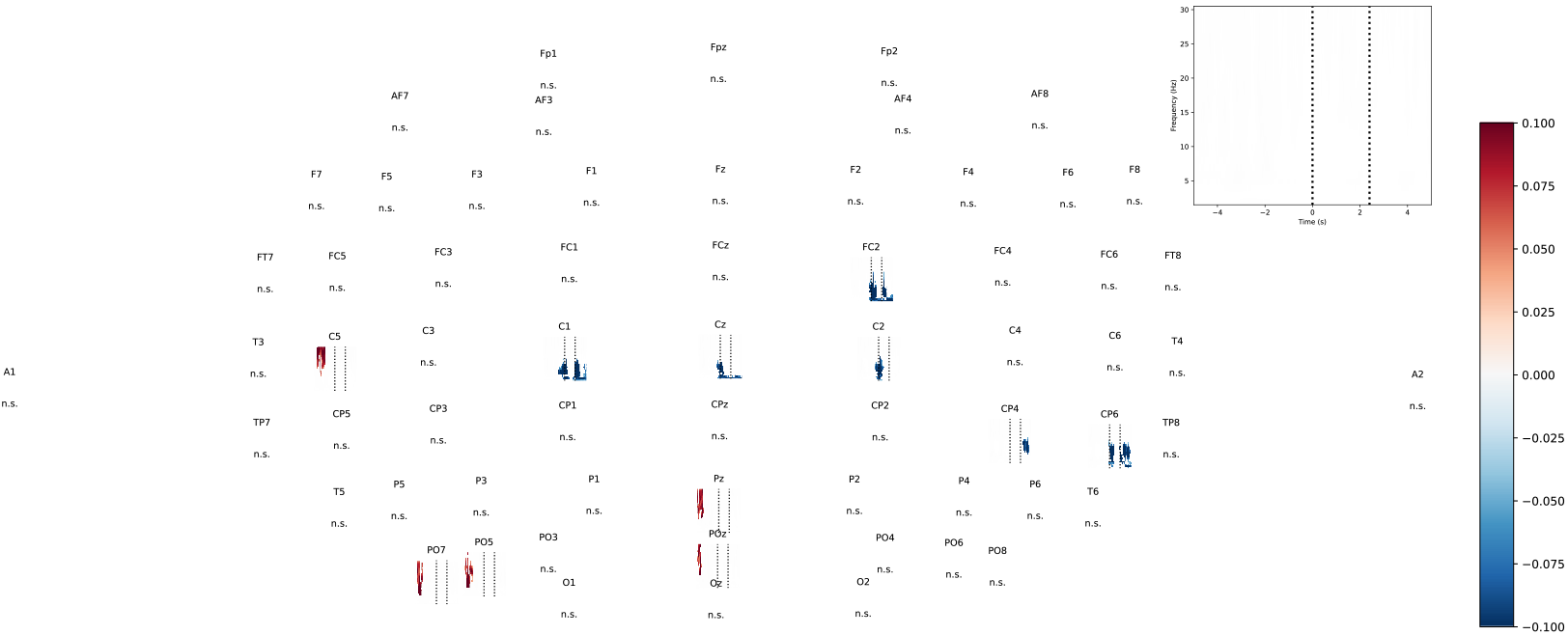
Results of the cluster-based permutation test for interoception group (high IAcc, low IAcc) differences in the familiar spectrogram. Only significant points are displayed. The dashed line at 0 s marks the onset of the silent interval, and the dashed line at 2.4 s indicates the resumption of the music. The upper-right inset shows the spectrogram axes (vertical frequency, horizontal time).

## Discussion

This study compared brain activity during a music recall task across familiarity groups (familiar, unfamiliar) and interoception groups (high IAcc, low IAcc). When comparing familiarity groups, the alpha-band spectrogram was more suppressed in the familiar group than in the unfamiliar group during melody listening and recall. As ERPs associated with music onset, N100 and P200 appeared in both familiarity groups. When examining ERPs at the onset of the silent interval, LPP was confirmed in the familiar group but was absent in the unfamiliar group. Furthermore, during music listening, SSEP was observed in both groups, whereas no SSEP-like peaks appeared during the silent interval. In comparing spectrograms across interoception groups, the high IAcc group showed a significant difference in alpha power between familiarity groups at the onset of silence, whereas the low IAcc group did not. A slight trend toward significance was also observed between interoception groups for the familiar group at the onset of silence.

### Spectrogram Comparison Between Familiarity Groups and Alpha-Power Suppression

Our experiment shows the results that the familiar stimulus group exhibited significantly lower power around 8–15 Hz (alpha and low-beta power) than the unfamiliar group in both music listening and recall. Notably, a significant difference emerged immediately after the start of the silent interval, at the onset of music recall. The previous research has reported that the alpha and low-beta power are significantly lower when participants listen to familiar music than to unfamiliar music [22] during the music listening task. Moreover, the power in the 8–14 Hz band has been found to be significantly lower for known music compared to unknown music, particularly right after the recall begins [25] during music recall.

In our study, the frequencies showing a significant difference between groups were in the 8–15 Hz range, aligning with these prior findings [22, 25]. The neural mechanisms underlying our results may be linked to the suppression of alpha power, which has been associated with attentional processes and expectation toward upcoming stimuli [40, 41], as well as with access to long-term memory [42]. In studies of word prediction, alpha power is suppressed more when the next word is easily predictable than when it is difficult to predict [43]. Based on such findings, it is possible that stronger attention is directed to a known melody during listening, and memories of the music are retrieved during recall.

The region of alpha-power suppression observed was at most electrodes, with especially pronounced suppression around the frontal and frontopolar regions. Similar suppression in these areas has been reported for music recall beyond the auditory cortex, including the inferior frontal cortex and medial frontal cortex [25] The inferior frontal cortex plays a role in melodic and pitch processing, as well as music recall [44], while the medial frontal cortex is involved in linking musical structure to memory [45]. The observed suppression of alpha power in these regions suggests that music retrieval engages complex cognitive processes, including memory retrieval and structural integration.

We used familiarity as an indicator of successful recall, consistent with previous research on music recall [29]. Other studies have similarly demonstrated that participants can recall familiar music when asked directly [46], reinforcing the validity of using familiarity as a recall measure. However, familiarity alone does not fully capture the vividness of recall. Incorporating a measure of recall vividness [47] could provide a more comprehensive assessment of recall success.

### Brain Activity Related to Music Listening and Imagery

During music listening, the N100 and P200 ERP components associated appeared to be familiar groups at most electrodes. For music recall, our result indicates that a peak resembling LPP was evoked in the familiar group of stimuli around the occipital and parietal areas. In contrast, no LLP component was observed for the unfamiliar group. Furthermore, both familiarity groups exhibited SSEPs at multiples of the 2.5 Hz tempo during music listening, but no SSEP-like peaks appeared during recall. These findings suggest that while the increase in spectrogram power upon music resumption likely originates from ERP responses.

From the results that LPP was only observed when a familiar melody was presented. The amplitude of the LPP has been reported to increase when a stimulus is associated with existing memories or knowledge [38, 48, 49]. This suggests that when music is successfully recalled, memory related to the music piece is actively retrieved. In contrast, when recall may be unsuccessful, access to or association with memory may be insufficient, causing the absence of the LPP component in an unfamiliar group.

Although SSEP was observed in both familiarity groups during music listening, neither group exhibited SSEP during the silent interval. Previous studies show that SSEP appears at frequencies matching the musical tempo during listening [24], as well as during the imagined rhythm of periodic stimuli [50]. Hence, the absence of SSEP during the silent interval in our result indicates that imagining a melody differs in neural processing from imagining a rhythm. The presence of music may make it more difficult to perceive the tempo and create more complexity in identifying the underlying rhythm, which could explain the absence of SSEP in silent intervals. This would require further exploration to understand how musical elements impact tempo perception.

### Influence of Interoception on Spectrogram During Music Recall

#### Relationship Between Alpha Power Suppression and Interoception in Music Recall

We compared the spectrograms between familiarity and interoceptive groups during silent intervals. As a result, alpha power tended to be more suppressed for familiar than unfamiliar music. This tendency was more pronounced in the high IAcc group. On the other hand, no significant differences were observed between interoceptive groups in the unfamiliar group. In addition, we did not observe significant differences between familiarities in the low IAcc group.

Alpha power has been reported to be significantly suppressed when attention is directed inward (e.g., interoceptive tasks) compared to rest, whereas it tends to increase in tasks focusing on external stimuli [27, 51]. These findings suggest that music recall is related to interoception. Moreover, if music recall involves directing attention inward, it may be characterized as directing attention to interoceptive processes. However, changes in alpha power when focusing on interoceptive signals differ depending on the specific experimental condition. For example, focusing on breathing can suppress alpha power [27], whereas focusing on heartbeat can increase alpha power [26]. The present study indicates that alpha power during music recall varies with the strength of interoception, but more investigation is needed to fully elucidate the effect of interoception on alpha power.

During tasks requiring attention to interoception, alpha power is significantly suppressed compared to resting conditions, whereas the tasks requiring attention to external stimuli are associated with increased alpha power [27, 51]. Given these findings, our alpha suppression results suggest that music recall may be related to interoceptive accuracy. However, the alpha power modulation during interoceptive attention has been reported to vary depending on the experimental conditions. For example, alpha power is suppressed when attention is focused on breathing [27]. In contrast, another study reported that it increases when attention is focused on heartbeat sensations [26]. The present study suggests that alpha power during musical recall varies depending on the intensity of interoceptive sensations. However, further discussion is needed to conclude the influence of interoception on alpha power.

It is important to note that we only compared participants with high vs. low interoceptive accuracy; the participants between these two extremes were excluded in this study. Thus, there remains the possibility that low rather than high interoceptive accuracy influences recall ability, which requires further investigation. In addition, the heartbeat counting task (HCT) used to assess interoceptive accuracy may reflect factors such as guessed heartbeat rates or imagined rhythms rather than true heartbeat perception [15, 52]. While participants were instructed not to rely on their usual heart rate or to physically detect their pulse [32], their internal sense of rhythm may still have influenced the results. Indeed, individuals with weaker interoception may be better at rhythm perception than those with stronger interoception [16]. Thus, the validity of using HCT as an index of interoceptive accuracy remains open to debate.

#### Elevated Beta Power During Music Listening and Attention to Interoception

From our experiment, we observed significantly higher beta power in the high IAcc group than in the low IAcc group during music listening. A positive correlation between beta power and IAcc has been reported for tasks involving attention to interoception. This finding suggests that the participants with higher IAcc may more effectively utilize interoceptive attention [51]. In this study, participants were instructed to imagine the melody in their minds, regardless of whether it was present or absent. It is possible that the participants with stronger interoception were simultaneously creating their own internal “playback,” distinct from the external auditory input.

#### Role of Insular Cortex in Music Recall

It has been reported that people with high interoceptive accuracy show increased activity in the insular cortex when listening to emotionally moving music [13]. The insular cortex is a key region for processing interoceptive signals [12]. Adopting the hypothesis that music recall involves directing attention toward interoception, the insular cortex could play a crucial role in the information flow underlying music recall. However, it is challenging to pinpoint deep-brain regions such as the insula using EEG. Future investigations employing imaging methods with higher spatial resolution may be required to elucidate the involvement of the insular cortex in music recall.

### Limitations and Future Directions

Given that the insular cortex processes interoceptive signals and that music recall appears to engage interoceptive attention, the insular cortex may play an important role in the information flow underlying music recall. However, deep-brain regions such as the insular cortex could not be directly evaluated using EEG. Moreover, the number of participants in this study was small; only participants with high or low interoceptive accuracy were included, leaving those in the middle range excluded from the analysis. Investigating participants in this middle range is an important direction for future research.

## Conclusion

This study investigated the effect of interoception on EEG activity during music recall. We compared spectrograms of music recall by familiarity (familiar vs. unfamiliar) and by interoception group. The results showed that alpha power suppression was more pronounced in the familiar group than in the unfamiliar group. This suppression was also stronger in participants with higher interoceptive accuracy compared to those with lower accuracy. Moreover, SSEP was observed during music listening; regardless of music familiarity, no SSEP peaks appeared during the silent interval. Additionally, an LPP was observed at the onset of the silent interval only in the familiar group. These findings suggest that music recall involves directing attention toward interoception.

## Supplementary Materials

**S1. Table Musical pieces stimuli**. The melody lists that were used as stimuli. Three 8-bar melodies were extracted from each of the 40 pieces, yielding 120 total stimuli.

**S2. Table HCT scores (IAcc)**. The score was calculated based on the standard formula for each participant.

**S3. Figure Time-frequency analysis results at 64 electrodes**. The grand-average difference in spectrograms between the familiarity results and their cluster-based permutation test results.

**S4. Figure ERP at the onset of the silent interval in familiar and unfamiliar groups**. The results show for all electrodes.

**S5. Table Statistical test results on N100 during the silent interval of the familiar music stimuli (***p <* 0.05 **highlighted in red)**

**S6. Table Statistical test results on N100 during the silent interval of the unfamiliar music stimuli (***p <* 0.05 **highlighted in red)**

**S7. Table Statistical test results on P200 during the silent interval of the familiar music stimuli (***p <* 0.05 **highlighted in red)**

**S8. Table Statistical test results on P200 during the silent interval of the unfamiliar music stimuli (***p <* 0.05 **highlighted in red)**

**S9. Table Statistical test results on P300 during the silent interval of the familiar music stimuli (***p <* 0.05 **highlighted in red)**

**S10. Table Statistical test results on P300 during the silent interval of the unfamiliar music stimuli (***p <* 0.05 **highlighted in red)**

**S11. Table Statistical test results on LPP during the silent interval of the familiar music stimuli (***p <* 0.05 **highlighted in red)**

**S12. Table Statistical test results on LPP during the silent interval of the unfamiliar music stimuli (***p <* 0.05 **highlighted in red)**

**S13. Figure ERP evoked at the melody resumption interval in the familiar and unfamiliar groups**

**S14. Table Statistical test results on N100 during the melody resumption of the familiar music stimuli**

**S15. Table Statistical test results on N100 during the melody resumption of the unfamiliar music stimuli (***p <* 0.05 **highlighted in red)**

**S16. Table Statistical test results on P200 during the melody resumption of the familiar music stimuli (***p <* 0.05 **highlighted in red)**

**S17. Table Statistical test results on P200 during the melody resumption of the unfamiliar music stimuli (***p <* 0.05 **highlighted in red)**

**S18. Table Statistical test results on P300 during the melody resumption of the familiar music stimuli (***p <* 0.05 **highlighted in red)**

**S19. Table Statistical test results on P300 during the melody resumption of the unfamiliar music stimuli (***p <* 0.05 **highlighted in red)**

**S20 Table Results of a two-way repeated-measures ANOVA on the PSD peaks at multiples of the melody tempo (2.5 Hz, 5.0 Hz, 7.5 Hz, 10 Hz)**. The independent variables were familiarity (familiar, unfamiliar) and interval (melody, silent).

**S21 Table Paired t-test comparisons of familiarity groups at each frequency**

**S22 Table Paired t-test comparisons of the interval factor at each frequency**

**S23 Table ANOVA results for each frequency band at the onset of the silent period (early silent period)**

**S24 Table ANOVA results for each frequency band during the late silent period**

**S25 Table Paired** *t***-test results for each frequency band at the onset of the silent period (early silent period)**

**S26 Table Paired** *t***-test results for each frequency band during the late silent period**

## Data Availability

All recorded data and stimuli used in the experiment are available in Zenodo (DOI: 10.5281/zenodo.15086711).

## Acknowledgments

This work was supported by JSPS Grant 21K18311. We would like to thank Editage (www.editage.jp) for English language editing.

---

**Table S1.** Melodies used as stimuli. Three 8-bar melodies were extracted from each of the 40 pieces, yielding 120 total stimuli.

| Composer | Title |
| --- | --- |
| L. v. Beethoven | Symphony “Ode to Joy” |
| A. Dvořák | Symphony No.9 “New World” |
| E. W. Elgar | Pomp and Circumstance |
| G. F. Handel | Messiah “Hallelujah” |
| G. Holst | Planets “Mercury” |
| F. Mendelssohn | Wedding March |
| W. A. Mozart | Eine Kleine Nachtmusik |
| W. A. Mozart | Turkish March |
| Hermann Necke | Csikos Post |
| J. Offenbach | Orpheus in Hell |
| P. I. Tchaikovsky | Swan Lake “Scene” |
| P. I. Tchaikovsky | The Nutcracker “March” |
| John Newton | Amazing Grace |
| The Sherman Brothers | It’s a Small World |
| J.L.Pierpont | Jingle Bells |
| popular English lullaby | Twinkle Twinkle Little Star |
| popular English lullaby | We Wish You a Merry Christmas |
| L. Harline | When You Wish upon a Star |
| American Folk Songs | Yankee Doodle |
| American Folk Songs | B-I-N-G-O |
| E. Satie | The Gymnopédies |
| I. Albeniz | Piano Sonate Op.82 |
| L. v. Beethoven | Piano Sonate Op.14–1 |
| A. Diabelli | Sonatine Op.151-2 |
| A. Dvořák | Waltz |
| A. Dvořák | Serenade for Strings Op. 22-5 |
| G. Faure | Dolly Suite, Kitty-valse |
| E. H. Grieg | Lyric Pieces |
| F. J. Haydn | Keyboard Sonata No.12 |
| F. J. Haydn | Keyboard Sonata No.28 |
| F. J. Haydn | Keyboard Sonata No.33 |
| F. Kuhla | Sonatine Op.55-1 |
| Theodor Leschetizky | Humoresque |
| F. Mendelssohn | Songs Without Words Op.19-1 |
| W. A. Mozart | Piano Sonate KV309 |
| S. S. Prokofiev | 10 Pieces Op.12–2 Gavotte |
| S. S. Prokofiev | 10 Pieces Op.12–3 Rigaudon |
| F. P. Schubert | Piano Sonata No.4 Scherzo |
| F. P. Schubert | Piano Sonata No.6 mov.3 |
| W. R. Wagner | Piano Sonata Op. 1 |

**Table S2.** HCT scores (IAcc) calculated based on the standard formula.

| Participants | IAcc |
| --- | --- |
| mk1p | 0.746484 |
| mk2p | 0.808750 |
| mk3p | 0.941536 |
| mk4p | 0.985581 |
| mk5p | 0.630218 |
| mk6p | 0.582161 |
| mk7p | 0.541054 |
| mk8p | 0.681867 |
| mk9p | 0.687332 |
| mk10p | 0.871105 |
| mk11p | 0.818032 |
| mk12p | 0.671805 |
| mk13p | 0.831785 |
| mk14p | 0.802572 |
| mk15p | 0.982950 |
| mk16p | 0.989899 |
| mk17p | 0.876810 |
| mk18 | 0.896149 |
| mk19 | 0.950631 |
| mk20 | 0.972434 |
| mk21 | 0.738436 |
| mk22 | 0.829377 |

**Table S20.** Results of a two-way repeated-measures ANOVA on the PSD peaks at multiples of the melody tempo (2.5 Hz, 5.0 Hz, 7.5 Hz, 10 Hz). The independent variables were familiarity (familiar, unfamiliar) and interval (melody, silent).

| Frequency | Factor | F Value | p Value | $\eta^2$ |
| --- | --- | --- | --- | --- |
| **2.5 Hz** | Familiarity (familiar, unfamiliar) | 19.3 | < 0.001 | 0.921 |
|  | Interval (melody, silent) | 76.2 | < 0.001 | 3.63 |
|  | Interaction | 2.08 | 0.1641 | 0.0990 |
| **5.0 Hz** | Familiarity (familiar, unfamiliar) | 16.1 | < 0.001 | 0.765 |
|  | Interval (melody, silent) | 176.0 | < 0.001 | 8.38 |
|  | Interaction | 17.2 | < 0.001 | 0.818 |
| **7.5 Hz** | Familiarity (familiar, unfamiliar) | 17.6 | < 0.001 | 0.838 |
|  | Interval (melody, silent) | 57.4 | < 0.001 | 2.73 |
|  | Interaction | 9.94 | < 0.01 | 0.473 |
| **10.0 Hz** | Familiarity (familiar, unfamiliar) | 0.311 | 0.583 | 0.0148 |
|  | Interval (melody, silent) | 43.7 | < 0.001 | 2.08 |
| | Interaction | $2.55 \times 10^{-4}$ | 0.9874 | $1.21 \times 10^{-5}$ |

**Fig S3.**
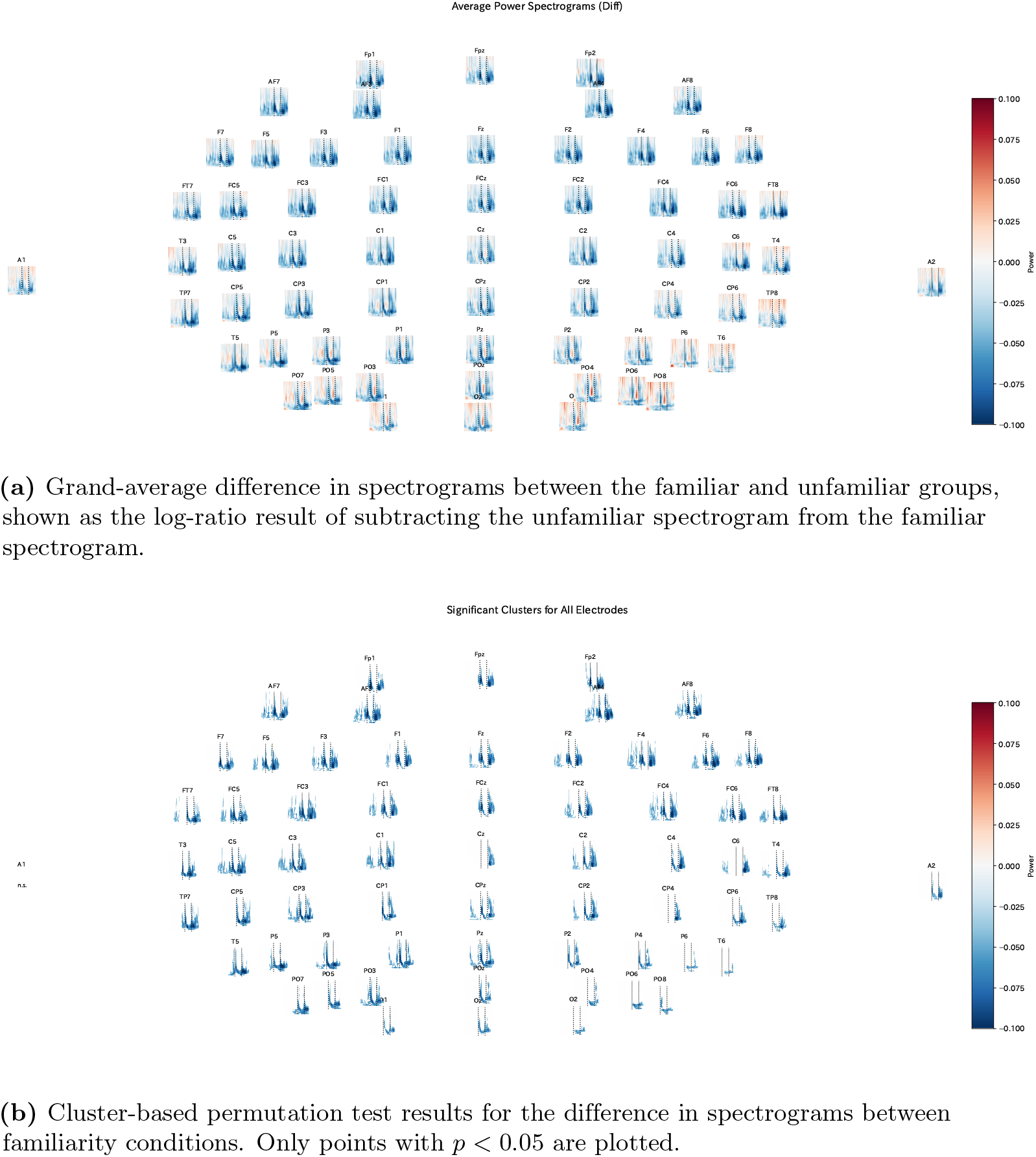
Time-frequency analysis results at 64 electrodes

**Table S21.** Paired t-test comparisons of familiarity groups at each frequency.

| Frequency | Comparison | $t$ Value | $p$ Value | Cohen's $d$ |
| --- | --- | --- | --- | --- |
| **2.5 Hz** | familiar (melody) vs. unfamiliar (melody) | 3.17 | 0.0228 | 0.677 |
|  | familiar (silent) vs. unfamiliar (silent) | 3.58 | 0.0105 | 0.764 |
| **5.0 Hz** | familiar (melody) vs. unfamiliar (melody) | -4.36 | < 0.01 | -0.93 |
|  | familiar (silent) vs. unfamiliar (silent) | 0.365 | 1.0 | 0.0778 |
| **7.5 Hz** | familiar (melody) vs. unfamiliar (melody) | 3.96 | < 0.01 | 0.844 |
|  | familiar (silent) vs. unfamiliar (silent) | 0.76 | 1.0 | 0.162 |
| **10.0 Hz** | familiar (melody) vs. unfamiliar (melody) | 0.338 | 1.0 | 0.072 |
|  | familiar (silent) vs. unfamiliar (silent) | 0.602 | 1.0 | 0.128 |

**Fig S4.**
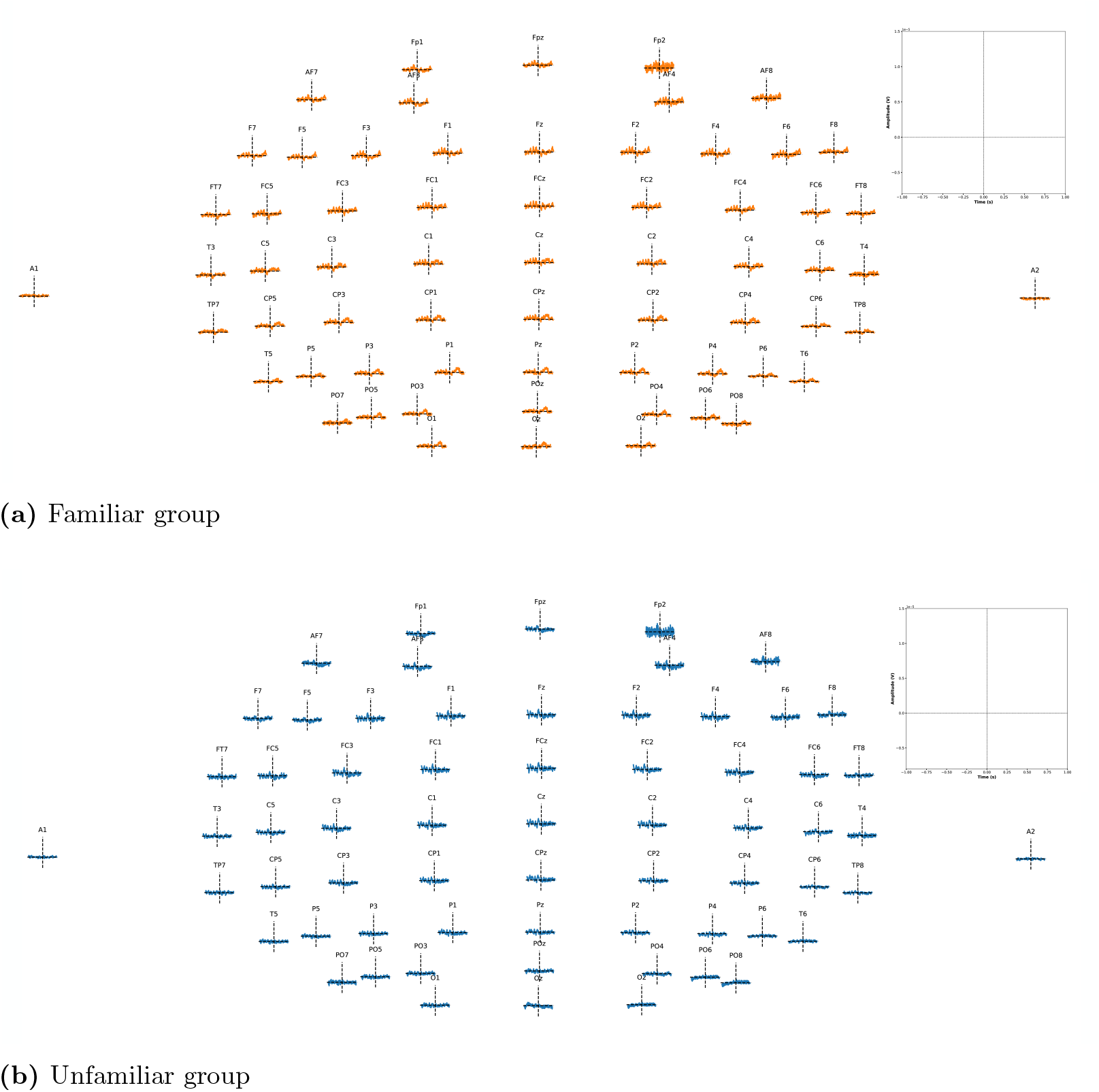
ERP at the onset of the silent interval for all electrodes. The dashed line on the horizontal axis denotes 0 *μ*V, and the dashed line on the vertical axis at 0 s indicates the start of the silent interval. The upper-right inset shows the axes (vertical amplitude, horizontal time).

**Table S22.** Paired t-test comparisons of the interval factor at each frequency.

| Frequency | Comparison | *t* Value | *p* Value | Cohen's *d* |
| --- | --- | --- | --- | --- |
| **2.5 Hz** | unfamiliar (melody) vs. unfamiliar (silent) | 7.85 | < 0.001 | 1.67 |
|  | familiar (melody) vs. familiar (silent) | 6.41 | < 0.001 | 1.37 |
| **5.0 Hz** | unfamiliar (melody) vs. unfamiliar (silent) | 13.3 | < 0.001 | 2.83 |
|  | familiar (melody) vs. familiar (silent) | 11.3 | < 0.001 | 2.42 |
| **7.5 Hz** | unfamiliar (melody) vs. unfamiliar (silent) | 5.18 | < 0.001 | 1.10 |
|  | familiar (melody) vs. familiar (silent) | 7.46 | < 0.001 | 1.59 |
| **10.0 Hz** | unfamiliar (melody) vs. unfamiliar (silent) | 4.88 | < 0.001 | 1.04 |
|  | familiar (melody) vs. familiar (silent) | 8.11 | < 0.001 | 1.73 |

**Fig S13.**
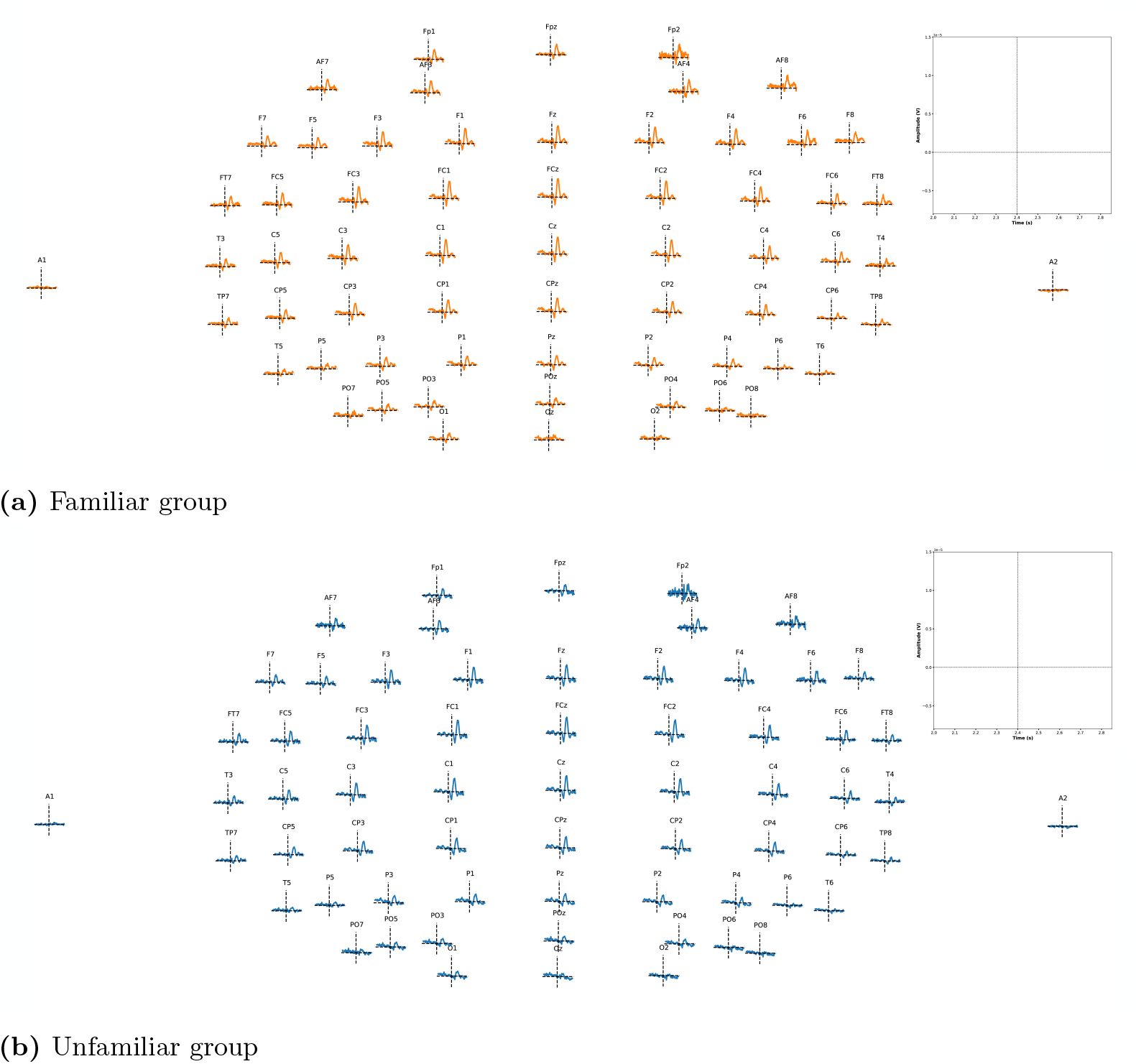
ERP evoked at the melody resumption interval in the familiar and unfamiliar groups. The dashed line on the horizontal axis denotes 0 *μ*V, and the dashed line on the vertical axis at 2.4 s indicates the point at which the melody resumes. The upper-right inset shows the axes (vertical amplitude, horizontal time).

**Table S23.** ANOVA results for each frequency band at the onset of the silent period (early silent period)

| Frequency | Factor | F Value | p Value | $\eta_p^2$ |
| --- | --- | --- | --- | --- |
| **Alpha (8-12Hz)** | Interception (between-subject effect) | 0.34 | 0.574 | 0.033 |
|  | Familiarity | 5.42 | 0.042 | 0.352 |
|  | Interaction | 11.75 | 0.006 | 0.540 |
| **Low Beta (13-16Hz)** | Interception (between-subject effect) | 0.005 | 0.946 | 0.0005 |
|  | Familiarity | 2.53 | 0.143 | 0.202 |
|  | Interaction | 7.31 | 0.022 | 0.422 |

**Table S24.**
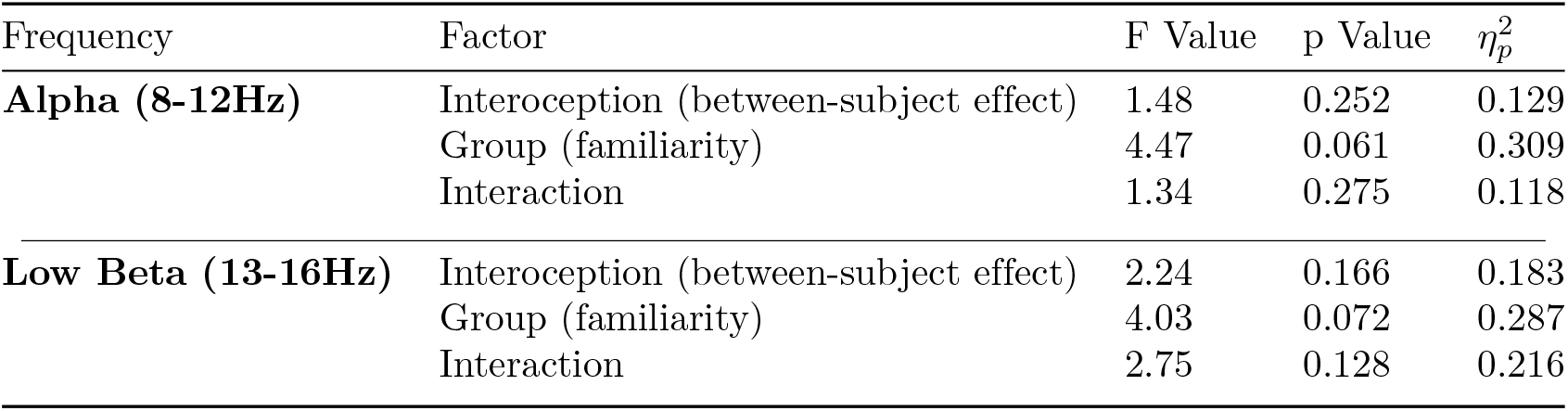
ANOVA results for each frequency band during the late silent period.

**Table S25.** Paired *t*-test results for each frequency band at the onset of the silent period (early silent period)

| Frequency Band | Comparison | *t* Value | *p* Value |
| --- | --- | --- | --- |
| **Alpha (8-12 Hz)** | high IAcc, familiar vs low IAcc, familiar | -2.386 | 0.117 |
|  | high IAcc, familiar vs high IAcc, unfamiliar | -4.071 | 0.010 |
|  | high IAcc, unfamiliar vs low IAcc, unfamiliar | 1.418 | 0.505 |
|  | low IAcc, familiar vs low IAcc, unfamiliar | 0.777 | 0.863 |
| **Low Beta (13-16 Hz)** | high IAcc, familiar vs low IAcc, familiar | -1.239 | 0.613 |
|  | high IAcc, familiar vs high IAcc, unfamiliar | -3.037 | 0.052 |
|  | high IAcc, unfamiliar vs low IAcc, unfamiliar | 1.115 | 0.687 |
|  | low IAcc, familiar vs low IAcc, unfamiliar | 0.786 | 0.859 |

**Table S26.** Paired *t*-test results for each frequency band during the late silent period1.

| Frequency Band | Comparison | *t* Value | *p* Value |
| --- | --- | --- | --- |
| **Alpha (8-12 Hz)** | high IAcc, familiar vs low IAcc, familiar | 0.320 | 0.988 |
|  | high IAcc, familiar vs high IAcc, unfamiliar | -2.313 | 0.160 |
|  | high IAcc, unfamiliar vs LI, unfamiliar | 1.661 | 0.372 |
|  | low IAcc, familiar vs low IAcc, unfamiliar | -0.679 | 0.903 |
| **Low Beta (13-16 Hz)** | high IAcc, familiar vs low IAcc, familiar | 0.337 | 0.986 |
|  | high IAcc, familiar vs high IAcc, unfamiliar | -2.593 | 0.104 |
|  | high IAcc, unfamiliar vs low IAcc, unfamiliar | 2.161 | 0.174 |
|  | low IAcc, familiar vs low IAcc, unfamiliar | -0.247 | 0.994 |

## Notes

### Competing Interest Statement

The authors have declared no competing interest.

